# Urinary Signatures Predict Calorie Restriction-Mediated Weight Loss in Obese Diversity Outbred Mice

**DOI:** 10.1101/2025.07.18.665483

**Authors:** Evan M. Paules, Isis Trujillo-Gonzalez, Melissa VerHague, Jody Albright, Delisha Stewart, Susan J. Sumner, Susan L. McRitchie, David Kirchner, Michael F. Coleman, Brian J. Bennett, Annie Green Howard, Penny Gordon-Larsen, John E. French, Stephen D. Hursting

**Author notes:** Corresponding Author: Stephen D. Hursting; Email:.

## Abstract

Predictive analytics encompassing metabolomic profiles are increasingly being used to forecast responders to dietary interventions. Advances using this approach are particularly needed to personalize and enhance the effectiveness of dietary weight loss interventions. Using obese Diversity Outbred (DO) mice that model genetic and phenotypic heterogeneity of human populations, we aimed to identify urinary metabolite signatures predictive of responsiveness to calorie restriction (CR)-mediated weight loss. DO mice (150 males, 150 females) were fed a high-fat diet for 12 weeks to induce obesity, then urine was collected and an 8-week CR regimen (30% decrease in energy intake) initiated. At study completion, mice were rank-ordered according to their percent body weight change, with mice in the extreme quartiles deemed CR responders (n=67) versus nonresponders (n=67). Targeted semi-quantitative metabolomics identified elevated glutamic acid and hydroxyproline as key urinary metabolites that distinguish CR responders from CR nonresponders, independent of sex. Three urinary metabolites (glutamic acid, hydroxyproline, and putrescine) distinguished male CR responders from nonresponders. Six metabolites (glutamic acid, hydroxyproline, dopamine, histamine, lysine, and spermine) distinguished female CR responders from nonresponders. Multivariate receiver operating characteristic analyses integrated the common metabolites and sex-specific metabolites to reveal moderate (males) to robust (females, males plus females) prediction models of CR-mediated weight loss. Further, pathway analysis identified several metabolic pathways, including arginine and proline metabolism, and alanine, aspartate, and glutamate biosynthesis, that distinguished CR responders from nonresponders and could be indicative of metabolic reprogramming to enhance insulin sensitivity and energy metabolism.

## Background

Defined as a body mass index ≥30 kg/m², obesity results from complex interactions among genetic, environmental, and behavioral factors (1–3). Obesity affects over 40% of U.S. adults (4, 5) and significantly raises the risk for chronic diseases, including type 2 diabetes, cardiovascular conditions, and multiple cancers (6). Efforts to reduce obesity prevalence are often hindered by individual variability in response to weight loss interventions (7, 8). Weight loss interventions tailored to an individual’s anthropometric data and psychosocial factors are more effective in supporting weight loss than generic interventions (9). Predictive analytics encompassing metabolomic profiles are increasingly being used to forecast responders to treatments in cardiovascular and other chronic diseases (10). Integrating metabolic predictors into weight loss interventions for obesity could enhance personalization and improve outcomes (11); however, studies investigating metabolic targets, especially in noninvasively collected biospecimens like urine, for predicting responses to weight loss interventions remain limited (12).

Calorie restriction (CR) is a cost-effective, non-pharmacological weight loss strategy, but achieving and maintaining significant weight loss through CR remains difficult for many (13). Biomarkers predictive of who will (or will not) favorably respond to CR-mediated weight loss are needed to facilitate more precise and sustainable anti-obesity treatment approaches (8). In a recent preclinical proof-of-principle study, we demonstrated that serum metabolic biomarkers including leptin can distinguish between obese Diversity Outbred (DO) mice that do, versus do not, respond favorably to CR-mediated weight loss (14). The DO mice are derived from the partially inbred Collaborative Cross Recombinant Lines, thus creating genetically unique individual mice and replicating the heterogeneity seen in human populations (15). Moreover, the DO mice exhibit extensive genetic diversity (∼45 million variants) and sex-specific metabolic traits, mirroring human responses to low-calorie diets (8, 16). Urine, as compared with serum, plasma and whole blood, offers several advantages as a biospecimen, including noninvasive sampling and ease of larger volume collection (17, 18). In this follow-up study, we investigated whether urinary metabolites could serve as biomarkers of CR responsiveness, hence extending our earlier serum-based findings (14) toward building a more comprehensive predictive signature for weight loss.

## Materials and Methods

### Animal diets and urine collection

All protocols were reviewed and approved by the Institutional Animal Care and Use Committee (IACUC protocol number: 21-003) at the North Carolina Research Campus in Kannapolis, NC. All animal care methods were previously described (14) and thus presented here in brief. Diversity Outbred (J:DO) mice from The Jackson Laboratory (stock #009376) were obtained at 8 weeks of age and were housed at 24°C with a 12-hour light/dark cycle and fed control diet (10 kcal% fat; Research Diets Catalog # D12450J) ad libitum for a 2 week acclimation period, then switched to a high-fat diet (60 kcal% fat; Research Diets #D12492) fed ad libitum for 12 weeks to elicit diet-induced obesity (DIO). Urine was collected using the Hatteras Chiller System (Grantsboro, NC, USA), with mice temporarily housed individually in cages with a wire mesh floor overnight following 12 weeks of the high-fat diet. Throughout the 24-hour collection period, urine samples were kept at 7°C, and after collection, samples were aliquoted and stored at -80°C until analysis. All mice were then housed in individual cages for the remainder of the study, fed the control diet ad libitum for 3 days, then administered a 30% CR diet (Catalog # F0078, Bio-Serv, Flemington, NJ) for 8 weeks. To achieve the CR regimen, all mice received a daily aliquot of food providing 70% of the 3-day average energy intake, and 100% of the vitamins, minerals, and amino acids, consumed during the control diet regimen. Following the 8 weeks of CR, all mice were fasted for 4 hours, following the start of the light cycle, and subsequently euthanized. Body weight was recorded weekly using a calibrated scale. Body composition, including fat mass and lean mass, was measured at baseline, after DIO, and following CR using an EchoMRI-100H analyzer (EchoMRI LLC, Houston, TX, USA). Humane endpoints throughout the study included any signs of significant distress (ruffled coat or hunched posture), disease, and weight loss that was greater than 20% within one week. During the 23 weeks of dietary intervention, 29 mice died spontaneously (20 males: 14 during DIO induction period, 6 during CR treatment period; 9 females: 6 during DIO induction period, 3 during CR treatment period). Mice that prematurely died were censored from subsequent analysis.

### Targeted semiquantified metabolomics analysis

Urine samples were received by the UNC Metabolomics and Exposome Laboratory for inventory and randomized using SAS version 9.4 (SAS, Cary, NC) prior to preparation for data acquisition. Urine was analyzed using a Biocrates Absolute*IDQ*® p180 kit (Biocrates, Life Sciences AG, Innsbruck, Austria) following the manufacturer’s instructions (16). In brief, internal standards, calibration standards (quantitation), quality control samples (interplate correlation and normalization), or 10 µL of experimental urine samples were placed at the center of each filter in the upper plate of a 96-well kit and dried with a nitrogen evaporator. Next, a 5% phenylisothiocyanate solution was added for derivatization of amino acids and biogenic amines. Following incubation, the filters were dried again with nitrogen. The metabolites were then extracted using a 5 mM ammonium acetate solution in methanol and transferred to a lower 96-deep well plate via centrifugation. Biocrates Absolute*IDQ*® p180 plates were analyzed using a AB Sciex API 4000 Triple Quad LC/MS (AB Sciex, Framingham, MA) equipped with an Agilent 1200 HPLC (Agilent, Santa Clara CA, USA). Amino acids and biogenic amines were determined in LC-MS mode. Quantification was carried out using internal standards and a calibration curve (Cal 1 to Cal 7). Data was managed on MetIDQ software (Biocrates, Life Sciences AG, Innsbruck, Austria), and all urinary metabolite data was normalized using creatinine (19).

### Data analysis

Data preprocessing was completed using MetIDQ Carbon software for peak extraction, automatic checking of peak alignment, normalization, and background deduction of mass spectrometry raw data. Orthogonal partial least squares discriminant analysis (OPLS-DA), receiver operating characteristic (ROC) curves, and pathway analyses were performed using MetaboAnalyst (20). Preprocessed data was uploaded to the MetaboAnalyst web server followed by a data integrity check with missing value imputations. All data was subsequently normalized using median values of each metabolite followed by log transformation and auto scaled.

Metabolites deemed important to the differentiation of study groups met the following thresholds determined using a fold-change analysis and t-test: fold change > 1.5 or < 0.66, and a raw *p* < 0.05. Metabolites corresponding to Variable Importance in Projection (VIP) values >1.50 (from the first principal component of the OPLS-DA analyses) were also deemed important to the differentiation of groups. Multivariate ROC curves were generated using OPLS-DA algorithm and predicted class probabilities were determined using 100 cross-validations. The area under the curve (AUC) was calculated to assess model discrimination. To guard against overfitting, model complexity was optimized to minimize validation errors, and permutation testing (n = 100) was performed to confirm that the observed AUC was not due to chance. Results are reported as mean AUC values with confidence intervals based on cross-validation replicates. All metabolite concentrations were subject to pathway analyses using the KEGG pathway in MetaboAnalyst. To identify pathways of interest, each pathway had to meet the following criteria: raw p-value <0.05 and an impact score of >0.3 (21).

## Results

### Body Mass and Composition of DO Mice Following High-Fat Diet and Calorie Restriction

Among 150 male and 150 female DO mice fed a high-fat diet for 12 weeks to induce obesity, then a 30% CR diet regimen for 8 weeks to induce weight loss (12; Fig. 1), significant heterogeneity in body mass and composition occurred in response to each diet for all mice, with similar variability by sex (Table 1). For all mice after the high-fat diet treatment, body weights ranged from 19.8 g to 70.7 g (median: 44.9 g in males, 35.9 g in females), percent fat mass from 11.2% to 55.0% (median: 34.8% in males, 39.0% in females), and percent lean mass from 40.8% to 88.1% (median: 60.1% in males, 56.8% in females). For all mice after CR treatment, body weight ranged from 15.8 g to 60.8 g (median: 35.4 g for males; 26.4 g for females), percent fat mass from 0.1% to 47.1% (median: 23.8% for males, 22.4% for females), and percent lean mass from 47.6% to 94.0% (median: 71.8% for males, 73.0% for females).

**Figure 1.**
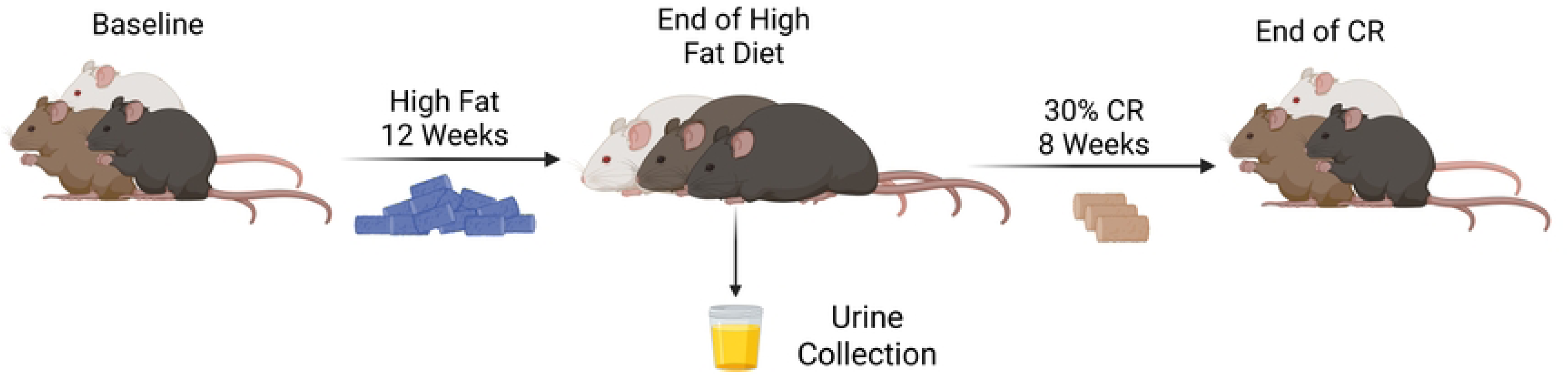
Timeline of the high-fat diet, urine collection, and calorie restriction.

**Table 1.** DO mice body weight and composition at baseline, following a high-fat diet, and following calorie restriction.

| <u>Intervention</u> | <u>Baseline</u> |  | <u>High-Fat Diet</u> |  | <u>Calorie Restriction</u> |  |
| --- | --- | --- | --- | --- | --- | --- |
|  | <u>Males and Females</u> |  |  |  |  |  |
| <u>Outputs</u> | <u>Median</u> | <u>Range</u> | <u>Median</u> | <u>Range</u> | <u>Median</u> | <u>Range</u> |
| Body Weight (g) | 21.4 | 13.5-33.8 | 39.1 | 19.8-70.7 | 30.4 | 15.8-60.8 |
| Fat Mass (%) | 12.7 | 3.6-33.0 | 36.0 | 11.2-55.0 | 23.1 | 0.1-47.1 |
| Lean Mass (%) | 83.1 | 62.5-95.0 | 59.0 | 40.8-88.1 | 71.9 | 47.6-94.0 |
|  | <u>Males</u> |  |  |  |  |  |
|  | <u>Median</u> | <u>Range</u> | <u>Median</u> | <u>Range</u> | <u>Median</u> | <u>Range</u> |
| Body Weight (g) | 23.9 | 17.6-33.8 | 44.9 | 25.2-70.7 | 35.4 | 20.0-60.8 |
| Fat Mass (%) | 10.8 | 3.6-21.6 | 34.8 | 11.2-50.4 | 23.8 | 0.1-44.4 |
| Lean Mass (%) | 85.6 | 72.7-95.0 | 60.1 | 43.5-88.1 | 71.8 | 50.3-91.5 |
|  | <u>Females</u> |  |  |  |  |  |
|  | <u>Median</u> | <u>Range</u> | <u>Median</u> | <u>Range</u> | <u>Median</u> | <u>Range</u> |
| Body Weight (g) | 19.9 | 13.5-27.1 | 35.9 | 19.8-57.0 | 26.4 | 15.8-46.6 |
| Percent Fat Mass (%) | 14.7 | 5.8-33.0 | 39.0 | 11.9-55.0 | 22.4 | 2.5-47.1 |
| Percent Lean Mass (%) | 80.4 | 62.5-88.8 | 56.8 | 40.8-82.0 | 73.0 | 47.6-94.0 |

Males and females: baseline: n=300, high-fat diet: n=280, calorie restriction: n=271; Males: baseline: n=150, high-fat diet: n=136, calorie restriction: n=130; Females: baseline: n=150, high-fat diet: n=144, calorie restriction: n=141

### Designation of CR Responders and Nonresponders by Percent Changes in Body Weight

We rank-ordered the mice (by both combined sex and separate sex) according to their CR-mediated percent weight change and then designated the mice in the extreme quartiles as CR nonresponders (quartile 1, n=67 mice with the least weight loss, including 32 males and 35 females) and CR responders (quartile 4, n=67 mice with the greatest weight loss, including 33 males and 34 females) (Table 2). Overall, the median percent change in body weight for CR responders was -38.6% (males: -38.0%; females -39.7%), compared to -8.7% for CR nonresponders (males: -7.4%, females: -12.1%).

**Table 2.** CR-induced percent body weight loss of CR responders and CR nonresponders.

**Change in Bodyweight (%)**
| CR nonresponders |  | CR responders |  |
| --- | --- | --- | --- |
| Median % Change | Range | Median % Change | Range |
| <u>**Males and Females**</u> |  |  |  |
| -8.7 (n=67) | -14.1 to 23.5 | -38.6 (n=67) | -57.0 to -31.8 |
| <u>**Males**</u> |  |  |  |
| -7.4 (n=32) | -11.9 to 23.5 | -38.0 (n=33) | -57.7 to -31.8 |
| <u>**Females**</u> |  |  |  |
| -12.1 (n=35) | -14.1 to 5.9 | -39.7 (n=34) | -57.0 to -35.0 |

### Predictive Urinary Metabolite Signatures of CR-Mediated Weight Loss in Obese Mice

To identify urinary metabolites that distinguish between CR responders and nonresponders, we performed targeted semiquantitative metabolomics and subsequent downstream data analyses on urine samples collected in obese DO mice immediately prior to CR intervention. First, we conducted a supervised OPLS-DA of urinary metabolites (Fig. 2A-C). Class separation, albeit often weak, was detected between CR responders and nonresponders in all mice (Q2=0.34, R^2^Y=0.48), males (Q2=0.07, R^2^Y=0.44), and females (Q2=0.43, R^2^Y=0.65).

**Figure 2.**
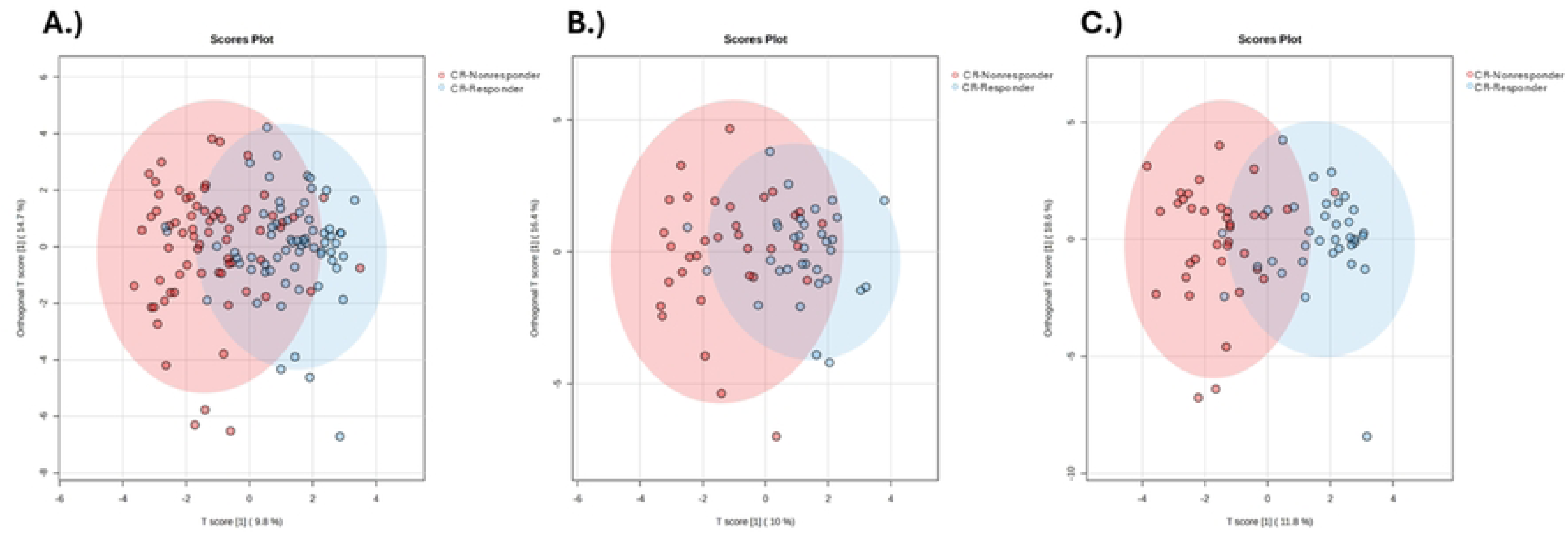
OPLS-DA Scores Plots of CR responder and CR nonresponder Urinary Metabolites. Scores plots of (A) all CR responders (blue circles) and nonresponders (red circles), (B) male CR responders and CR nonresponders, and (C) female CR responders and CR nonresponders.

Next, using the OPLS-DA statistics, we identified urinary metabolites that had a VIP value >1.50. We also determined metabolites (comparing CR responders versus CR nonresponders) with a fold change >1.5 or <0.66 and with *p* < 0.05. The following metabolites met these criteria: hydroxyproline and glutamic acid, in both sexes; hydroxyproline, glutamic acid, and putrescine in males; and hydroxyproline, glutamic acid, dopamine, histamine, lysine, and spermine in females (Table 3). The metabolites were consistently elevated in CR responders versus nonresponders, with the exceptions of putrescine, histamine, and spermine, which were suppressed. In multivariate ROC analyses, each of these 3 metabolic signatures, when applied to their associated sex-combined or sex-specific group, was at least moderately good (based on area under the curve, AUC, >0.70) at predicting CR responsiveness (Fig. 3A-C). Specifically, AUCs (95% CI) were 0.76 (0.69-0.86) in all mice combined, 0.71 (0.52-0.86) in males, and 0.85 (0.69-0.97) in females.

**Figure 3.**
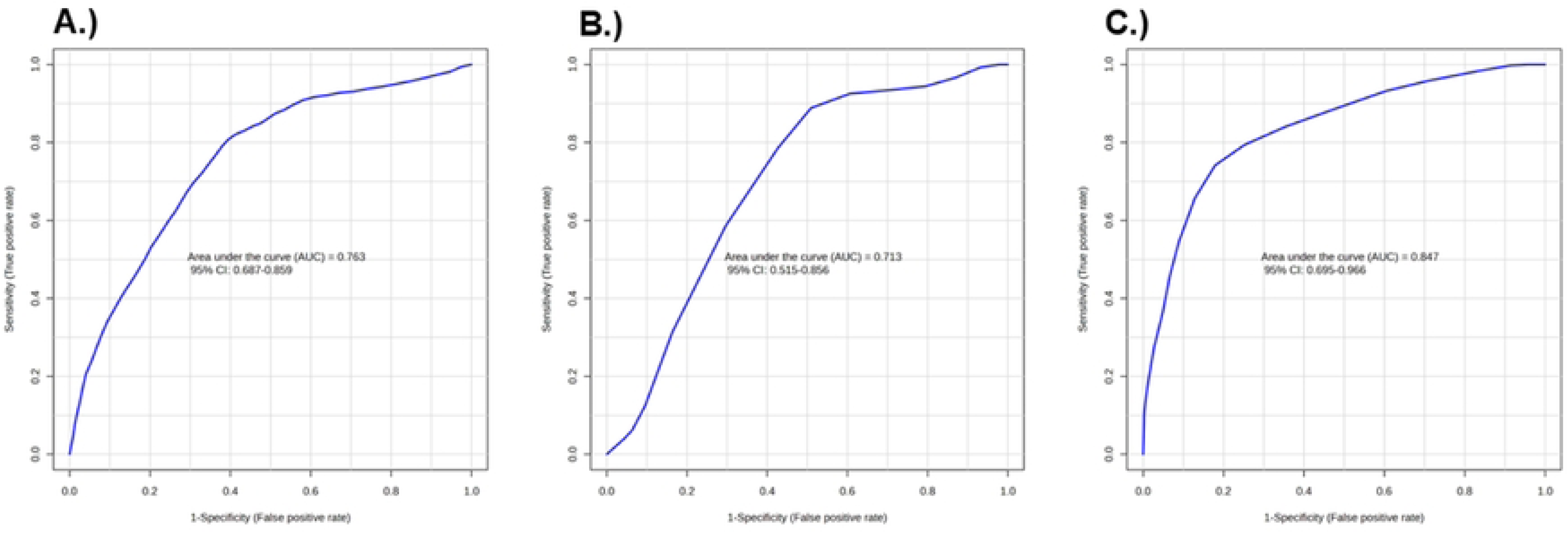
Multivariate ROC curves that distinguish CR responders and CR nonresponders Receiver operating characteristic (ROC) curves of (A) all CR responders and nonresponders (AUC: 0.76, confidence interval (CI): 0.68-0.85), (B) male CR responders and CR nonresponders (AUC: 0.71, CI: 0.51-0.85), and (C) female CR responders and CR nonresponders (AUC: 0.84, CI: 0.69-0.96).

**Table 3.** All, male, and female CR Responder vs CR Nonresponder. Top differential urinary metabolites.

| Metabolite | VIP | Fold Change | Raw p-value | Pathway | CR Responder vs Nonresponder |
| --- | --- | --- | --- | --- | --- |
| **<u>All</u>** |  |  |  |  |  |
| **Hydroxyproline** | 2.05 | 2.83 | <0.001 | Arginine and proline metabolism | ↑ |
| **Dopamine** | 1.90 | 1.74 | <0.001 | Catecholamine biosynthesis | ↑ |
| **Glutamic acid** | 1.92 | 3.12 | <0.001 | Arginine and proline metabolism | ↑ |
| **<u>Male</u>** |  |  |  |  |  |
| **Hydroxyproline** | 2.12 | 2.60 | 0.006 | Arginine and proline metabolism | ↑ |
| **Glutamic acid** | 1.92 | 3.81 | 0.001 | Arginine and proline metabolism | ↑ |
| **Putrescine** | 1.86 | 0.53 | 0.001 | Glutathione metabolism | ↓ |
| **<u>Female</u>** |  |  |  |  |  |
| **Dopamine** | 1.87 | 2.06 | <0.001 | Catecholamine biosynthesis | ↑ |
| **Hydroxyproline** | 1.75 | 3.23 | <0.001 | Arginine and proline metabolism | ↑ |
| **Glutamic acid** | 1.64 | 2.23 | <0.001 | Arginine and proline metabolism | ↑ |
| **Histamine** | 1.63 | 0.39 | <0.001 | Histidine metabolism | ↓ |
| **Lysine** | 1.57 | 1.79 | <0.001 | Lysine degradation | ↑ |
| **Spermine** | 1.54 | 0.11 | <0.001 | Arginine and proline metabolism | ↓ |

### Pathway Topology Analysis of Urinary Metabolite Infers Differences in Metabolic State Between CR Responders and Nonresponders

We used pathway topology analysis of urinary metabolites to interrogate differences in the metabolic state of CR responders and nonresponders (Fig. 4A-C). Top pathways with *p* < 0.05 and pathway score > 0.3 were deemed to have high impact, and these included arginine and proline metabolism; histidine metabolism; glycine, serine, and threonine metabolism; and alanine, aspartate, and glutamate biosynthesis in all mice, males and females (Table 4). Other high-impact pathways identified were tyrosine metabolism, as well as phenylalanine, tyrosine, and tryptophan biosynthesis, in all mice and female mice; and taurine and hypotaurine metabolism in females.

**Figure 4.**
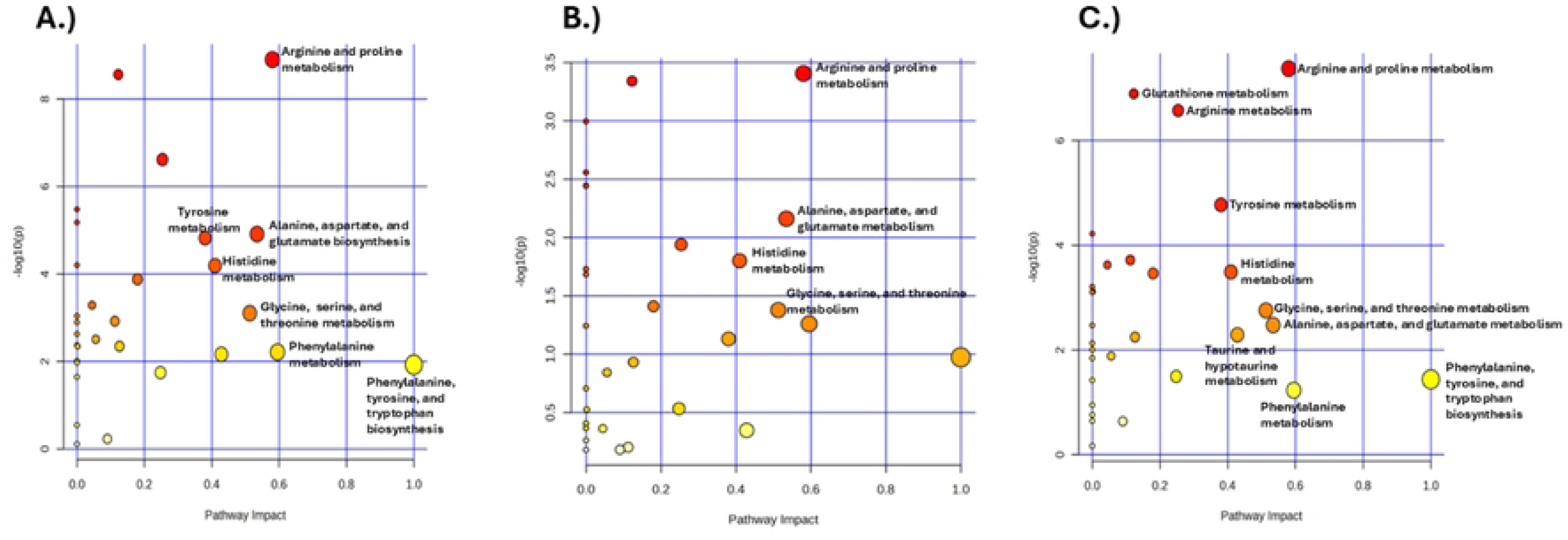
Pathway analysis differentiating CR responders and CR nonresponders urinary profiles Bubble chart of KEGG pathway analysis for (A) all, (B) male, and (C) female CR responders vs CR nonresponders. Top pathways with *p* < 0.05 and pathway score > 0.3 are labeled accordingly. Pathway impact values are on the x-axis while -log(p) values are on the y-axis.

**Table 4.** All, male, female top KEGG metabolic pathways that distinguish CR responders from CR nonresponders.

| <u>Pathway</u> | <u>Total</u> | <u>Hits</u> | <u>Raw p</u> | <u>Impact</u> |
| --- | --- | --- | --- | --- |
| <u>All</u> |  |  |  |  |
| Phenylalanine, tyrosine, and tryptophan biosynthesis | 4 | 2 | <0.01 | 1.0 |
| Arginine and proline metabolism | 36 | 8 | <0.001 | 0.58 |
| Tyrosine metabolism | 42 | 3 | <0.001 | 0.38 |
| Histidine metabolism | 16 | 4 | <0.001 | 0.40 |
| Glycine, serine, and threonine metabolism | 34 | 3 | <0.001 | 0.51 |
| Phenylalanine metabolism | 10 | 3 | <0.01 | 0.59 |
| Alanine, aspartate, and glutamate biosynthesis | 28 | 6 | <0.001 | 0.53 |
| <u>Males</u> |  |  |  |  |
| Arginine and proline metabolism | 36 | 8 | <0.001 | 0.58 |
| Histidine metabolism | 16 | 4 | <0.05 | 0.40 |
| Glycine, serine, and threonine metabolism | 34 | 3 | <0.05 | 0.51 |
| Alanine, aspartate, and glutamate biosynthesis | 28 | 6 | <0.01 | 0.53 |
| <u>Females</u> |  |  |  |  |
| Phenylalanine, tyrosine, and tryptophan biosynthesis | 4 | 2 | <0.05 | 1.0 |
| Arginine and proline metabolism | 36 | 8 | <0.001 | 0.58 |
| Tyrosine metabolism | 42 | 3 | <0.001 | 0.38 |
| Histidine metabolism | 16 | 4 | <0.001 | 0.40 |
| Glycine, serine, and threonine metabolism | 34 | 3 | <0.01 | 0.51 |
| Alanine, aspartate, and glutamate biosynthesis | 28 | 6 | <0.01 | 0.53 |
| Taurine and hypotaurine metabolism | 8 | 1 | <0.01 | 0.42 |

## Discussion

Using targeted semiquantitative metabolomics and downstream data analyses of biobanked urine samples from obese DO mice (150 males, 150 females) collected immediately before CR intervention (14), we identified urinary metabolic signatures that distinguished CR-mediated weight loss responders from nonresponders with good discrimination (multivariate ROC curves, AUC > 0.70). The metabolic signatures included hydroxyproline and glutamic acid for all the mice; hydroxyproline, glutamic acid, and putrescine for the males; and hydroxyproline, glutamic acid, spermine, dopamine, histamine, and lysine for the females. Further, we discerned high-impact metabolic pathways, including arginine and proline metabolism and alanine, aspartate, and glutamate biosynthesis, that distinguished CR responders from nonresponders and could indicate metabolic reprogramming to enhance insulin sensitivity and energy metabolism. Hence, these preclinical data i) demonstrate that metabolites in the noninvasive biofluid urine can forecast CR-mediated weight loss, and ii) complement and extend the previously reported CR weight-loss predictive ability of metabolites in systemic circulation (14).

Of the urinary metabolites assessed, only hydroxyproline and glutamic acid distinguished CR responders from CR nonresponders in each combined- and separate-sex group. Each of these metabolites was also identified as being part of the arginine and proline metabolic pathway, as determined by KEGG pathway analysis. Arginine and proline metabolism primarily takes place in the intestinal-renal axis (22). Independent of protein translation, arginine is an important intermediary metabolite in the urea cycle and polyamine biosynthesis, while proline is involved in collagen biosynthesis (23). Glutamic acid concentrations impact the metabolism of arginine since glutamic acid produces α-ketoglutarate, which can be converted into aspartate and, subsequently, arginine. Putrescine (differentiating the male groups) and spermine (differentiating the female groups) are also involved in arginine and proline metabolism, likely due to arginine’s role in polyamine biosynthesis. Additionally, three distinct metabolic pathways—histidine metabolism; alanine, aspartate, and glutamate biosynthesis; and glycine, serine, and threonine metabolism—differentiated CR responders from CR nonresponders in all mice combined. Several urinary metabolites linked to CR response are part of these pathways. For example, histamine and glutamic acid are part of the histidine metabolism pathway, while lysine plays a role in glycine, serine, and threonine metabolism.

The literature supports both glutamic acid and hydroxyproline as indicators of metabolic distress (22–28). Urinary glutamic acid levels are considered a reliable proxy for circulating levels (24). Moreover, increased circulating glutamic acid in human studies directly correlate with obesity and indicate possible metabolic acidosis (25). Elevated glutamic acid may also suggest reduced liver functionality in CR responders. As metabolic dysfunction-associated steatohepatitis progresses, there is a shift in glutamine extraction and catabolism to glutamic acid from the liver to the kidneys to compensate for the metabolic acidosis and dysregulated liver metabolism (26, 27). Hydroxyproline is one of the most prevalent amino acids in collagen (28). In a healthy state, hydroxyproline is a substrate for extracellular matrix production, which helps to preserve the integrity and functionality of hepatocytes (28). However, hepatic stellate cells activate upon chronic hepatocyte damage, leading to their increased proliferation, excessive production of extracellular matrix (28, 29) and initiation of liver fibrosis (28). In humans and rodents, circulating and urinary hydroxyproline levels are correlated with liver disease, specifically, the presence and severity of fibrosis (28), and may be a result of degradation of newly synthesized collagen, particularly in the beginning stages of fibrosis (30).

Spermine and putrescine are polyamines that play important roles in cellular metabolism and physiology (31). High polyamine intake is linked to lower all-cause mortality in a prospective community-based cohort study (32), and elevated polyamine levels are linked to enhanced energy metabolism and increased autophagy (31). Additionally, polyamines function as free radical scavengers to reduce the harmful effects of reactive oxygen species (33). However, elevated excretion of polyamines has been shown to be associated with chronic disease and inflammation. Moreover, enhanced catabolism of cellular polyamines can also increase cellular oxidative stress (34). In our study, spermine and putrescine, respectively, were reduced in female and male CR responders, compared to CR nonresponders, respectively. Putrescine can be converted into spermine as well as back-converted into putrescine through spermidine synthase. Spermidine synthase is regulated, in part, by androgens (35) and could possibly explain the sexual dimorphism identified in male and female CR responders.

Dopamine is a neurotransmitter involved in the regulation of energy balance via food intake (36). In rodents, the blocking of dopamine receptors results in mitigating the reward stimulus of food, which suggests that altered dopaminergic signaling contributes to the feeding patterns in individuals with obesity (37). In humans, women with obesity shown pictures of high-calorie foods had significantly greater activation in several regions of the brain that relate to reward processing including the amygdala and hippocampus, among others (38). Dopamine crosses the blood brain barrier through monoamine transporters and is excreted in the urine (39). We observed that urinary dopamine concentrations were significantly higher in female CR responders compared to female CR nonresponders, potentially related to how dopamine impacts eating behavior and reward responses. Moreover, dopamine was the top metabolite in our VIP analysis (Table 3). When food intake is restricted, CR responders may be unable to sustain dopamine-driven reward signaling, possibly altering hypothalamic circuits regulating satiety and energy expenditure. This is supported by evidence in humans, where higher urinary dopamine concentrations are associated with increased energy intake (40). However, in our preclinical study, urinary dopamine concentrations only distinguished female (and not male) CR responders from CR nonresponders, possibly due to estrogen regulation of dopamine release (41).

Histamine is a biogenic amine that regulates inflammation and is produced by a variety of immune cells, such as mast cells, as well as the glomeruli of the kidneys (42, 43). Urinary and plasma histamine levels are commonly increased in response to high levels of cytokines and chemokines as often occurs in allergic reactions (42, 44). High histamine levels in the urine, as observed in the CR nonresponders relative to CR responders may also indicate the development of chronic kidney disease, especially with the infiltration of mast cells into the kidneys being correlated with the development of nephropathy (42). However, the separation of CR responders and CR nonresponders by histamine levels was only identified in female mice and not male mice, possibly due in part to differential effects of testosterone versus estrogen on renal fibrosis (45).

Lysine is an essential amino acid obtained through metabolism of dietary proteins. The literature is mixed on lysine having a protective effect on kidney function in models of obesity and kidney disease or as a marker of obesity and diabetes. We identified that lysine, specifically as part of lysine degradation, was elevated in female CR responders compared to CR nonresponders. The lysine degradation pathways include the saccharopine pathway, predominately active in the kidney, and the pipecolate pathway, predominately active in the brain (46). In the short-to mid-term, elevated lysine is associated with protective effects in kidney disease in rodent models, yet the mechanisms underpinning these effects are unknown (46). Moreover, a lysine metabolite, α-aminoadipic acid, was found to be associated with diabetes risk and markers of obesity in humans (47).

Study limitations include a lack of 1) quantitation of physical activity, which is likely to be heterogeneous in these mice and may have impacted the amount of weight lost during CR; 2) data on estrous cycle in females, which may have influenced sex-steroid and other hormonal levels that can influence appetite and weight loss response. Moreover, we did not track water intake or determine glomerular filtration rate, which would have identified differences in kidney function that may have impacted urinary concentrations of certain urine metabolites. Future studies that account for these possible covariates and include similar analyses using urine samples from weight loss studies in humans are needed to extend these initial findings and ensure a robust metabolic signature predicting responsiveness to CR-mediated weight loss.

## Conclusions

Leveraging urine as a noninvasive biospecimen and using Diversity Outbred mice as a model of the phenotypic heterogeneity in human populations, we identified a urinary metabolite signature predictive of favorable CR-induced weight loss in obese mice. This metabolite signature and associated metabolic pathways, including arginine and proline metabolism, and alanine, aspartate, and glutamate biosynthesis, indicated CR responders, relative to nonresponders, were predisposed to exhibit greater weight loss, increased insulin sensitivity, and enhanced metabolic reprogramming in response to CR. Future work should focus on validating and extending the metabolic predictors of weight loss reported herein as well as integrating them with other predictors of weight loss response, such as serum leptin levels (14), to further personalize weight loss regimens and increase their success.

## Dataset Availability

Datasets analyzed for this study are available in UNC Dataverse at: https://doi.org/10.15139/S3/SJQLFJ

## Author Contributions

EMP analyzed and interpreted data, prepared graphics, and wrote the manuscript; ITG interpreted data and wrote the manuscript; MV, JA and MFC performed the animal experiments; DS, SLM and DK performed and analyzed metabolomic analysis; SJS provided funding and supervision for metabolomic analysis; AGH advised on the statistical analysis and reviewed the manuscript; JEF, PGL, and BJB conceived the study, obtained funding and interpreted data; SDH conceived the study, obtained the funding, interpreted data, and wrote the manuscript.

## List of Abbreviation

DO: Diversity outbred
CR: Calorie restriction
ROC: Reciver operating characteristic
OPLS-DA: Orthoganal partial least squares discriminant analysis
VIP: Variable importance projection
AUC: Area under the curve

## Acknowledgements

BioRender was used to create Figure 1.

(Created in BioRender. Paules, E. (2025) https://BioRender.com/d33s798)

